# KV7 but not dual SK/IK channel openers inhibit the activation of colonic afferents by noxious stimuli

**DOI:** 10.1101/2023.06.08.544183

**Authors:** Charity N Bhebhe, James P Higham, Rohit A Gupta, Tim Raine, David C Bulmer

**Author notes:** Author for correspondence: David C Bulmer.

## Abstract

In numerous subtypes of central and peripheral neurons, small and intermediate conductance Ca^2+^-activated K^+^ (SK and IK, respectively) channels are important regulators of neuronal excitability. Transcripts encoding SK channel subunits, as well as the closely related IK subunit, are co-expressed in the soma of colonic afferent neurons with receptors for the algogenic mediators adenosine triphosphate (ATP) and bradykinin, P2X_3_ and B_2_, highlighting the potential utility of these channels as drug targets for the treatment of abdominal pain in gastrointestinal diseases such as irritable bowel syndrome. Despite this, pre-treatment with the dual SK/IK channel opener SKA-31 had no effect on the colonic afferent response to ATP, bradykinin or noxious ramp distention of the colon. Inhibition of SK or IK channels with apamin or TRAM-34, respectively, yielded no change in spontaneous baseline afferent activity, indicating these channels are not tonically active. In contrast to its lack of effect in electrophysiological experiments, comparable concentrations of SKA-31 abolished ongoing peristaltic activity in the colon *ex vivo*. Treatment with the K_V_7 channel opener retigabine blunted the colonic afferent response to all applied stimuli. Our data therefore highlight the potential utility of K_V_7, but not SK/IK, channel openers as analgesic agents for the treatment of abdominal pain.

## Introduction

Ca^2+^-activated K^+^ (K_Ca_) channels couple intracellular Ca^2+^ signals to K^+^ efflux, thereby regulating membrane excitability in nerve and muscle, and cell volume and K^+^ homeostasis in non-excitable cells. The α subunits of these channels are split into three subclasses based on single channel conductance: SK1-3 (4-14 pS) encoded by *Kcnn1*-*3* (Köhler *et al*., 1996), IK (30-40 pS) encoded by *Kcnn4* (Ishii *et al*., 1997), and BK (200-300 pS) encoded by *Kcnma1* (Butler *et al*., 1993). Members of the K_Ca_ channel family also differ in their Ca^2+^ sensitivity, coupling to different Ca^2+^ sources, voltage-dependence (only BK is voltage-dependent) and pharmacological properties. SK channel α subunits have distinct but overlapping tissue expression, with functional channels formed of α subunit tetramers which can assemble as both homomers and heteromers, providing greater diversity within this family of K^+^ channels (Monaghan *et al*., 2004; Church *et al*., 2015).

SK channel subunits are expressed in peripheral sensory neurons (Usoskin *et al*., 2015; Zeisel *et al*., 2018) and there is now evidence for these channels playing a role in the processing of both innocuous and noxious stimuli. Immunofluorescent staining has shown co-expression of SK1 and SK2 channels with peptidergic and non-peptidergic markers in small-diameter sensory neurons in rat dorsal root ganglia (DRG), while SK3 is more broadly expressed (Mongan *et al*., 2005). Sensory input to the spinal cord during innocuous and noxious mechanical stimulation was elevated by inhibition of SK channels, while SK channel activation attenuated sensory input (Bahia *et al*., 2005). IK channels are also expressed in small-diameter sensory neurons (Mongan *et al*., 2005). What’s more, genetic ablation or pharmacological blockade of IK channels potentiated nocifensive behaviour following formalin or capsaicin administration, but had no effect on inflammatory or neuropathic pain (Lu *et al*., 2017).

Sensory signalling from the distal colon and rectum is mediated by the lumbar splanchnic and pelvic nerves and is of vital importance for defecation and the sensation of bloating, urgency and pain (Brierley, Hibberd and Spencer, 2018). A greater understanding of the ion channels which regulate colonic afferent activity will provide insights into the mechanisms underpinning visceral sensation and may highlight potential targets for future therapeutics. Whether SK/IK channels regulate colonic afferent function is currently unknown.

In this study, we have used pharmacological tools to compare the effect of SK/IK and K_V_7 channel activation on the colonic afferent response to algogenic stimuli and noxious distention of the colon. Our data demonstrate that pre-treatment with the SK/IK channel opener SKA-31 at concentrations which abolished peristaltic activity in the colon had no effect on the colonic afferent response to ATP, bradykinin, or noxious distention of the colon. In alignment with a previous report, subsequent activation of K_V_7 channels by retigabine robustly inhibited the colonic afferent response to each stimulus (Peiris *et al*., 2017). These data highlight the potential utility of K_V_7, but not SK/IK, channel openers for the treatment of abdominal pain.

## Methods and materials

### Animals

Experiments were performed using tissue from male C57Bl6 mice (10-14 weeks of age) euthanatized by exposure to rising concentrations of CO2 followed by cervical dislocation in accordance with Schedule 1 of the Animals (Scientific Procedures) Act 1986 Amendment Regulations 2012 and local ethical review by the University of Cambridge Animal Welfare and Ethical Review Body (AWERB). Mice were housed in cages of up to five littermates under a 12-hour light/dark cycle with enrichment (e.g., igloos and tunnels) and *ad libitum* access to food and water.

### In silico analysis of SK/IK transcript expression in colon projecting sensory afferents

To establish the expression of transcripts encoding SK and IK channel subunits, as well as receptors for ATP and bradykinin, we used a previously published, publicly available RNAseq dataset comprising back-labelled colonic sensory neurons (Hockley *et al*., 2019). Transcript expression is given as the base ten logarithm of transcripts per million. The proportion of colonic afferents co-expressing given transcripts was ascertained manually using conditional counting functions in Microsoft Excel.

### Colonic afferent fibre studies

Following euthanasia, the colorectum (from splenic flexure to anus) with the associated lumbar splanchnic nerve was isolated, removed and luminal content gently flushed. The tissue was transferred to a recording bath where it was cannulated to allow luminal perfusion against an end pressure of 2-5 mmHg (200 µL/min) and serosally superfused (7 mL/min; 32-34°C) with Krebs buffer (in mM: 124 NaCl, 4.8 KCl, 1.3 NaH_2_PO_4_, 25 NaHCO_3_, 1.2 MgSO_4_, 11.1 D-glucose and 2.5 CaCl_2_) supplemented with atropine (10 µM) and nifedipine (10 µM) to block smooth muscle contractility. Ongoing nerve discharge was recorded from isolated lumbar splanchnic nerve bundles (rostral to the inferior mesenteric ganglia) using borosilicate glass suction electrodes. Signals were amplified (gain, 5k), band-pass filtered (100-1500 Hz, Neurolog, Digitimer Ltd, UK) and digitally filtered for 50 Hz noise (Humbug, Quest Scientific, Canada) before being digitised (20 kHz, Micro1401, Cambridge Electronic Design, UK) and recorded using Spike2 software and displayed in a chart recorder format (Cambridge Electronic Design, UK). Nerve activity was quantified by counting the number of action potentials (spikes) passing a threshold set at a magnitude of twice the background noise and displayed as a rate histogram using the data analysis function in Spike2. Luminal pressure was recorded using a pressure transducer (Neurolog NL108) filled with Krebs buffer, signals were digitised at 100 Hz and displayed alongside nerve activity using Spike 2.

Changes in the colonic afferent response to ATP, bradykinin, and ramp distension (0-80 mmHg; performed by occluding the luminal outflow) were determined in separate tissues in the presence of pre-treatment with the SK/IK channel opener SKA-31 (100 µM) and the K_V_7 channel opener retigabine (100 µM), or respective vehicle control (DMSO 0.1%). For studies with ATP, colonic afferent responses were measured to three separate applications of ATP (20ml, 3mM) given 60 min apart. The second application of ATP was given in the presence of SKA-31 or vehicle and was preceded by a 50 mL pre-treatment with SKA-31 or vehicle. The third application of ATP was given in the presence of retigabine or vehicle and was preceded by a 50 mL pre-treatment with retigabine or vehicle. For studies with bradykinin, colonic afferent responses were measured to four separate applications of bradykinin (20 mL, 1 µM) given 30 min apart. The third application of bradykinin was given in the presence of SKA-31 or vehicle and was preceded by a 50 mL pre-treatment with SKA-31 or vehicle. The fourth application of ATP was given in the presence of retigabine or vehicle and was preceded by a 50 mL pre-treatment with retigabine or vehicle. For ramp distensions, the colonic afferent response to distension (0-80 mmHg, evoked by 200 µL/min luminal perfusion with Krebs buffer) were measured in response to seven separate distensions given 15 min apart. The fourth ramp distension was performed in the presence of SKA-31 or vehicle and was preceded by a 7 min pre-treatment with SKA-31 or vehicle. The sixth ramp distension was performed in the presence of retigabine or vehicle and was preceded by a 7 min pre-treatment with retigabine or vehicle. In addition, the effect of vehicle (50 mL, 0.1% DMSO), TRAM-34 (50 mL, 10 µM), apamin (50 mL, 1µM) on ongoing nerve discharge was examined in separate experiments.

### Data analysis

For tissues stimulated with either ATP or bradykinin, colonic afferent response curves were generated by subtracting baseline nerve activity, calculated as the mean nerve discharged over the 3 min period prior to administration of ATP or BK, from the ongoing nerve discharge measured in 1 min intervals from 6 mins prior to administration of ATP or BK to 24 mins afterwards (30 min in total). From these curves, the response to repeated ATP or BK challenges was measured as total spike firing (expressed as the area under afferent response curve from time 0-1440 s calculated using the trapezoidal method) and was statistically compared to vehicle treated preparations using a repeated measures one-way ANOVA with Holm-Sidak post-hoc test to determine the reproducibility of responses. Following this, the effect of drug pre-treatment was examined by comparing the calculated total spike firing (AUC) and mean changes in afferent activity for time matched preparations pre-treated with SKA-31 vs vehicle and retigabine vs vehicle using unpaired Students t-test for comparison of AUCs and two-way ANOVA for response profiles over time.

For tissue stimulated by ramp distension, colonic afferent response curves were generated by subtracting baseline nerve activity, calculated as the mean nerve activity over the 3 min period prior to luminal distension, from the ongoing nerve discharge measured in 5 mmHg intervals from 0-80 mmHg. From these curves, the effect of respective drug pre-treatment (SKA-31 or retigabine) was statistically compared with vehicle treated preparations using a two-way ANOVA, and the peak response to colorectal distension at 80 mmHg following drug pre-treatment (normalised to the peak response obtained from the distention prior to treatment) statistically compared with respective time matched vehicle pre-treatment using an unpaired Student’s t-test.

Changes in ongoing nerve discharge were also examined following pre-treatment with the IK channel blocker, TRAM-34, the SK channel blocker, apamin, and compared with respective changes in ongoing nerve discharge following pre-treatment with vehicle. The effect of SKA-31 or retigabine on ongoing nerve discharge (prior to treatment with bradykinin or ATP) was also investigated. Changes in ongoing nerve discharge were generated by subtracting baseline nerve activity, calculated as the mean nerve discharged over the 3 min period prior to administration of respective treatments, from the ongoing nerve discharge measured in 1 min intervals from 3 mins prior to administration to 6 mins afterwards. Changes were statistically compared between respective vehicle and treatment groups using a two-way ANOVA with Holm-Sidak post-hoc tests between time points.

### Colonic motility

Following euthanasia, the descending colon and rectum was removed and cannulated in a tissue bath superfused with Krebs buffer (7 mL/min; 32-34°C) and luminally perfused with Krebs buffer in an open system at a rate of 200 µL/min. Luminal pressure was monitored using a Neurolog NL108 pressure transducer, with the signals digitised (100 Hz, Micro1401, Cambridge Electronic Design, UK) and recorded using Spike2 (Cambridge Electronic Design, UK). Luminal pressure was raised to 5-10 mmHg by elevation of the outflow to trigger colonic migrating motor complexes (CMMCs). Tissue stabilised for at least 30 minutes before drug application (SKA-31, 70 mL, 100 µM). The amplitude of CMMCs in each experiment was determined by averaging the peak change in pressure for all individual CMMCs before and after drug application. The average CMMC frequency (per minute) before and after drug application was also determined. Changes in the amplitude and frequency of CMMCs were statistically compared before and after drug treatment using a paired Student’s t-test.

### Data analysis and statistics

All graphing and analysis was carried out in Prism 9 (GraphPad Inc.). Data were scrutinized to ensure they met the assumptions of parametric analyses and, where appropriate, non-parametric rank-based analyses were used. Sample sizes were not predetermined before data acquisition, but inter-group comparisons were decided before experiments began. P-value cut offs are displayed in figures as *p<0.05, **p<0.01, ***p<0.001, ****p<0.0001.

## Results

### Expression of SK channels in colonic afferent neurons

To ascertain the expression of transcripts encoding SK channel genes, we used previously published single cell RNAseq data from back-labelled murine colonic afferent neurons (Hockley *et al*., 2019). The expression of *Kcnn1-4* was examined across all colonic afferent subtypes (Figure 1A). While *Kcnn1-2* were broadly expressed in all colonic afferent subtypes, *Kcnn3* was poorly expressed (Figure 1A). *Kcnn4* was also expressed in colonic afferent neurons, though not to the same extent as *Kcnn1* and *Kcnn2* and was primarily expressed in non-peptidergic afferents. We also examined the expression of transcripts encoding receptors for ATP (P2X_3_, *P2rx3*) and bradykinin (B_2_ receptor, *Bdkrb2*) as these algogenic stimuli will be used later in this study. *P2rx3* was widely expressed across colonic afferent subtypes, while *Bdkrb2* was more selectively expressed in peptidergic afferents (Figure 1A).

**Figure 1:**
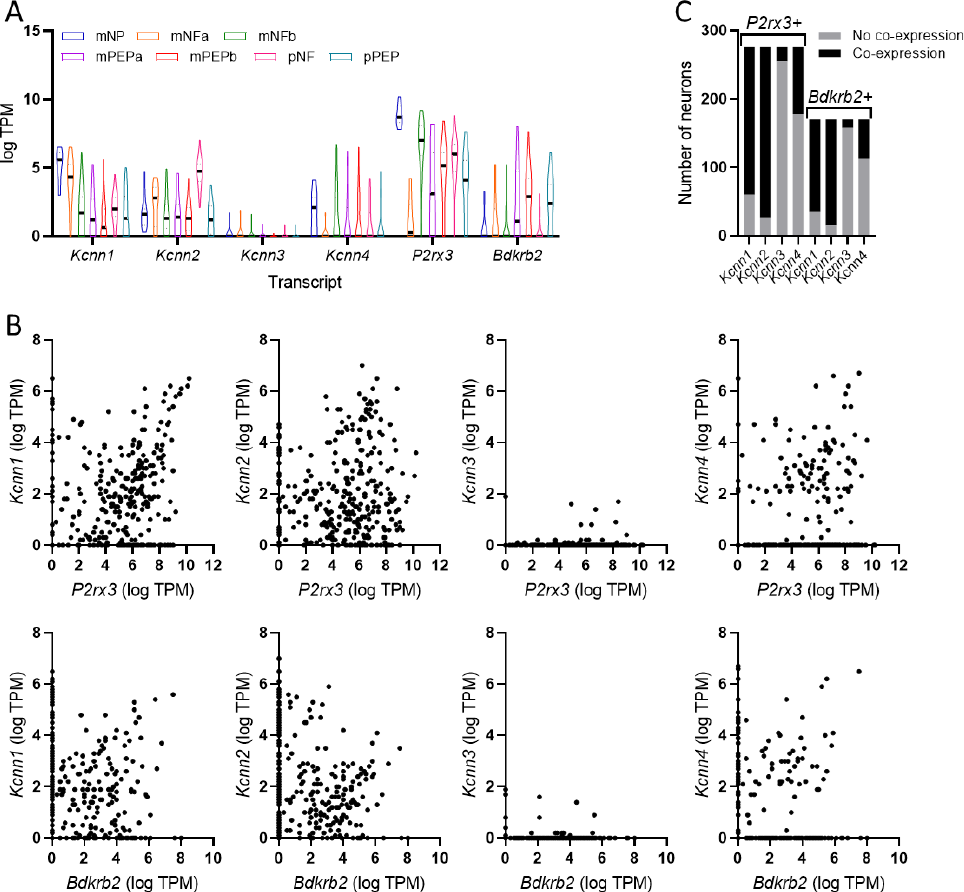
Expression of transcripts encoding SK and IK channel subunits in colonic sensory neurons. (A) Expression (expressed as transcripts per million, TPM) of transcripts encoding SK channel subunits and receptors for ATP (*P2rx3*) and bradykinin (*Bdkrb2*) across colonic sensory neuron subtypes. Neuronal subtypes are denoted: NP, non-peptidergic; NFa/b, neurofilament-expressing; PEP, peptidergic; m, mixed lumbar splanchnic and pelvic afferents; p, pelvic afferents. (B) Scatter plots showing the co-expression of *P2rx3* (top row) and *Bdkrb2* (bottom row) with *Kcnn1*-*4*. Each point represents a single colonic afferent neuron. (C) Grouped data showing the number of neurons expressing either *P2rx3* or *Bdkrb2* which co-express *Kcnn1*-*4*. Data redrawn from Hockley *et al*., 2019.

There was notable co-expression of *Kcnn1*-*2*, but less so of *Kcnn3*-*4*, with *P2rx3* and *Bdkrb2* (Figure 1B). 276 colonic afferent neurons expressed *P2rx3*, of which 216 (78.3%) and 250 (90.6%) co-expressed *Kcnn1* and *Kcnn2*, respectively (Figure 1C). *Bdkrb2* was expressed in 170 colonic afferent neurons, of which 135 (79.4%) and 155 (91.2%) co-expressed *Kcnn1* and *Kcnn2*, respectively (Figure 1C). *Kcnn3* was co-expressed in 7.6% and 7.1% of *P2rx3*- and *Bdkrb2*-expressing colonic afferent neurons, respectively (Figure 1C). *Kcnn4* was co-expressed in 35.5% and 33.5% of *P2rx3*- and *Bdkrb2*-expressing colonic sensory neurons, respectively (Figure 1C).

While P2X_3_ is the dominant ionotropic ATP receptor expressed by colonic sensory neurons, metabotropic ATP receptors (e.g., P2Y_1_) are also expressed by these afferents (Hockley *et al*., 2016, 2019). Of 98 neurons expressing *P2ry1* (encoding P2Y_1_), 81 (82.6%), 91 (92.3%), 7 (7.1%) and 35 (35.7%) co-expressed *Kcnn1*, *2*, *3* and *4*, respectively.

### SK/IK channels do not regulate the afferent response to ATP

Given the robust co-expression of *P2rx3* and *Bdkrb2* with transcripts encoding SK/IK channel subunits in colonic afferents, we used whole-nerve suction electrode recording of the lumbar splanchnic nerve (LSN) innervating the distal colon to test whether these channels regulate the afferent response to ATP and bradykinin. Bath application of ATP (3 mM) evoked a marked increase in afferent activity which did not desensitise and was not affected by co-application of DMSO (vehicle for later experiments, Figure 2Ai). There was no difference in the peak afferent activity (p = 0.96), or the total spikes discharged (p = 0.67, N = 5, Figure 2B) between the first, second and third applications of ATP.

**Figure 2:**
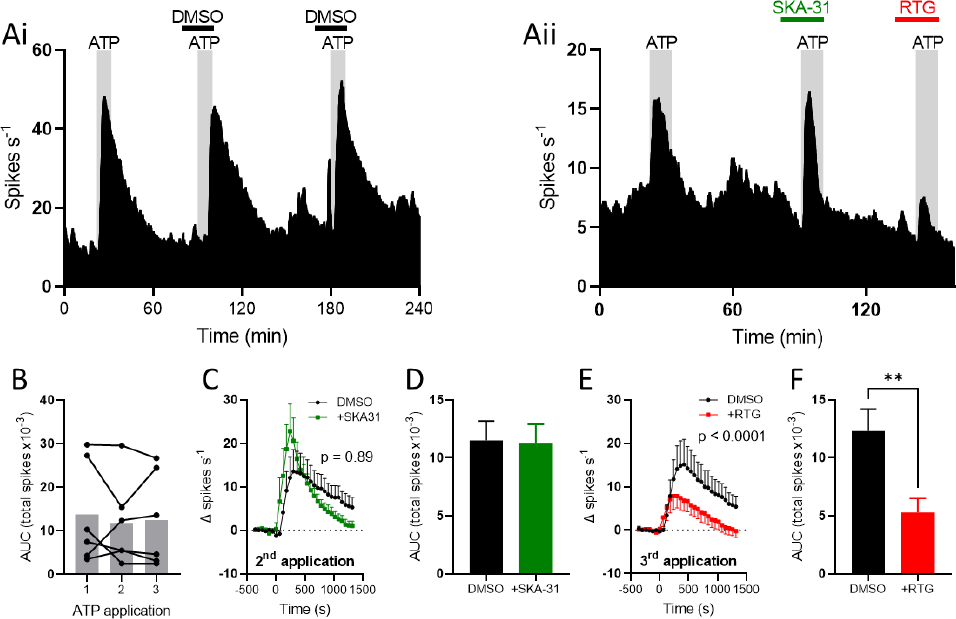
SK/IK channels do not regulate the afferent response to ATP. (A) Exemplar rate histograms showing afferent firing during repeated applications of ATP (*i*) with SKA-31 (*ii*) retigabine (RTG, *ii*). (B) Grouped data showing the total spike discharge following ATP application for each of the three applications. Bar shows mean, filled circles show individual experiments. One-way repeated measures ANOVA with Holm-Sidak post-tests. (C) Grouped data showing the change in afferent firing rate after the second ATP application (applied at 0 s) in DMSO-(black) and SKA-31-(green) treated tissue. Two-way ANOVA. (D) Grouped data showing the total spike discharge (area under curves in (C)) following the second ATP application. Two-tailed unpaired t-test. (E) Grouped data showing the change in afferent firing rate after the third ATP application (applied at 0 s) in DMSO- and retigabine-(red) treated tissue. Two-way ANOVA. (F) Grouped data showing the total spike discharge following the third ATP application. Two-tailed unpaired t-test.

SKA-31 (100 µM), an activator of SK (pEC_50_ = 5.5-5.7) and IK (pEC_50_ = 6.6) channels (Sankaranarayanan *et al*., 2009), had no affect the afferent response to ATP (Figure 2Aii). Afferent firing following the second ATP application was no different between DMSO- and SKA-31-treated tissues (main effect of drug, p = 0.89, Figure 2C). Total spike discharge was unaffected by treatment with SKA-31 (p = 0.94, N = 6) compared to DMSO-treated control tissue (N = 6, Figure 2D).

Retigabine (100 µM), an activator of K_V_7 channels, was co-applied with the third ATP application, resulting in a robust inhibition of the afferent response to ATP (Figure 2Aii). Afferent firing after ATP application was markedly reduced by retigabine compared to DMSO-treated tissue (main effect of drug, p < 0.0001, Figure E). Total spike discharge in control preparations was 12289±1889 spikes (N = 6), compared to 5321±1241 spikes in retigabine-treated preparations (p = 0.0067, N = 8, Figure 2F).

### SK/IK channels do not regulate the afferent response to bradykinin

Bath application of bradykinin (1 µM) gave rise to elevated afferent activity which underwent desensitisation with repeated applications (Figure 3A, *left*). The first application of bradykinin evoked greater afferent firing than all subsequent applications (p < 0.043), but there was no difference in afferent firing evoked by the second, third and fourth applications (p > 0.48, N = 5, Figure 3B). SKA-31 (100 µM) was applied with the third application of bradykinin (Figure 3A, *right*). Compared to DMSO-treated tissues, SKA-31 treatment did not affect afferent firing following bradykinin application (main effect of drug, p = 0.85, Figure 3C). Total afferent firing was very similar between treatment groups (6782±1180 spikes vs 6672±1401 spikes, p > 0.99, N = 5 per group, Figure 3D). Unlike SKA-31, retigabine pre-treatment before the fourth bradykinin application inhibited the response to bradykinin (Figure 3A, *right*). Bradykinin-evoked afferent firing was reduced in retigabine-treated compared to DMSO-treated tissues (main effect of drug, p = 0.033, Figure 3E). Total afferent discharge after bradykinin application was 6372±1014 spikes (N = 5) in DMSO-treated tissue and 1075±441 spikes (N = 6) in retigabine-treated tissue (p = 0.0005, Figure 3F). During the third application of bradykinin, afferent firing rate increased by 83.6±11.4% and 78.5±15.5% in DMSO- and SKA-31-treated tissues, respectively (p = 0.80, Figure 3G). During the fourth application of bradykinin, afferent firing increased by 76.2±5.7% and 24.3±4.1% in DMSO- and retigabine-treated tissues, respectively (p < 0.0001, Figure 3G).

**Figure 3.**
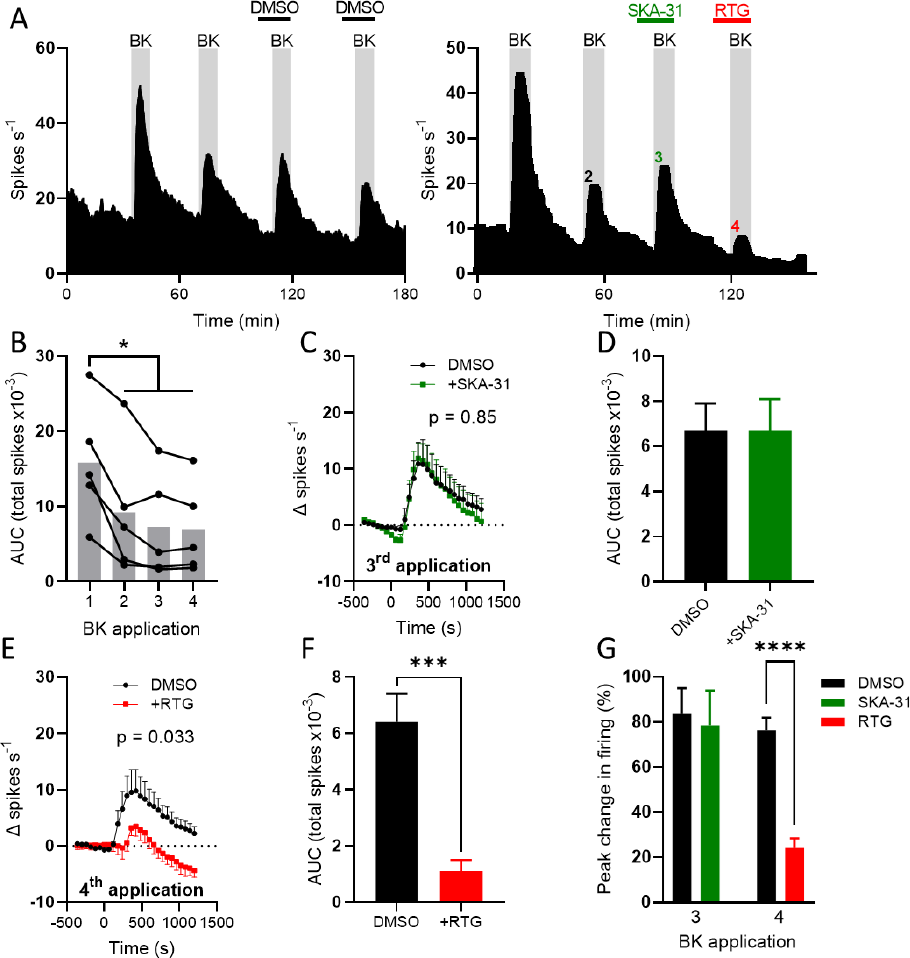
SKA-31 had no effect on bradykinin-evoked colonic afferent activity. (A) Exemplar rate histograms showing afferent firing during repeated applications of bradykinin (BK, *left*) with SKA-31 and retigabine (*right*). (B) Grouped data showing the total spike discharge following four successive applications of bradykinin. Bar shows mean, filled circles show individual experiments. One-way repeated measures ANOVA with Holm-Sidak post-tests. (C) Grouped data showing the change in afferent firing rate following the third application of bradykinin (applied at 0 s) in DMSO- and SKA-treated tissue. Two-way ANOVA. (D) Grouped data showing the total spike discharge following the third application of bradykinin. Two-tailed unpaired t-test. (E) Grouped data showing the change in afferent firing rate following the fourth application of bradykinin (applied a 0 s) in DMSO- and retigabine-treated tissue. Effect of retigabine on spontaneous activity subtracted from trace for time < 0 s. Two-way ANOVA. (F) Grouped data showing the total spike discharge following the fourth application of bradykinin. Two-tailed unpaired t-test. (G) Grouped data showing the percentage increase in afferent firing rate following the third and fourth application of bradykinin. Two-tailed unpaired t-test (third and fourth applications analysed separately).

### SK/IK channels do not regulate the afferent response to ramp distention of the colon

Ramp distention of the colon to a luminal pressure of ∼80 mmHg resulted in repeatable increases in afferent discharge (Figure 4A). Application of SKA-31 (100 µM) before and during the 4^th^ ramp distention had no effect on pressure-induced afferent firing compared to the preceding ramp distention (Figure 4A) or to DMSO-treated tissue (main effect of drug, p = 0.13, Figure 4B). Application of retigabine before and during the 6^th^ ramp distention reduced pressure-induced afferent firing compared to the preceding ramp distention (Figure 4A) and to DMSO-treated tissue (main effect of drug, p < 0.0001, Figure 4C). Retigabine inhibited afferent firing at luminal pressures >5 mmHg. During the 4^th^ ramp distention, afferent activity increased 88.5±3.4% (N = 6) and 94.7±7.7% (N = 6) in DMSO- and SKA-31-treated preparations, respectively (p = 0.48, Figure 4D). During the 6^th^ ramp distention, afferent activity increased 101.0±3.7% (N = 6) and 33.4±2.9% (N = 5) in DMSO- and retigabine-treated preparations, respectively (p < 0.0001, Figure 4D).

**Figure 4:**
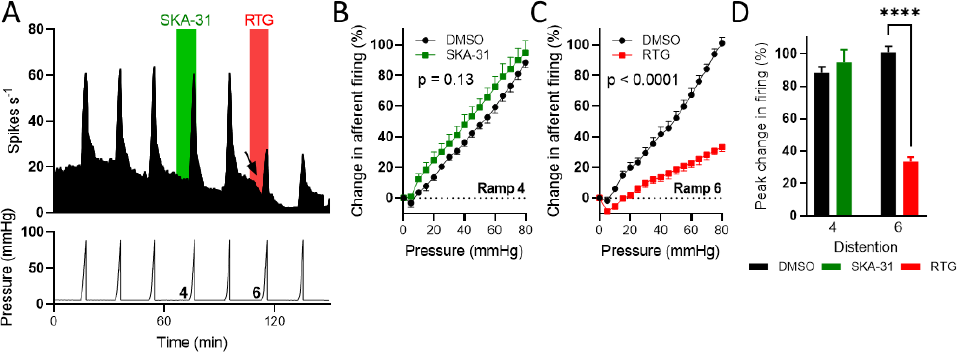
SKA-31 did not affect the afferent response to ramp distention of the colon. (A) Exemplar rate histogram (*top*) showing the afferent firing during repeated ramp distention of the colon to ∼80 mmHg luminal pressure (*bottom*). The application of SKA-31 and retigabine is shown in green and red, respectively. The arrow highlights the decrease in spontaneous afferent firing following retigabine application. (B) Pressure-response relationship for the fourth ramp distention in DMSO- and SKA-31-treated tissue. Two-way ANOVA. (C) Pressure-response relationship for the sixth ramp distention in DMSO- and retigabine-treated tissue. Two-way ANOVA. (D) Grouped data showing the percentage increase in afferent firing rate during the fourth and sixth ramp distention. Two-tailed unpaired t-test (fourth and sixth ramp distention analysed separately).

### SK/IK channels do not regulate spontaneous colonic afferent firing

We next investigated whether SK/IK channels tonically regulated spontaneous colonic afferent firing by recording ongoing afferent activity before and after the application of either apamin (SK channel blocker, 1 µM) or TRAM-34 (IK channel blocker, 10 µM). Application of apamin failed to affect afferent activity compared to DMSO application (main effect of drug, p = 0.55, Figure 5A). Afferent firing rate was decreased by 0.83±0.33 spikes s^−1^ (N = 4) and 0.89±0.45 spikes s^−1^ (N = 4) following the application of DMSO and apamin, respectively (p = 0.94, Figure 5B). Similarly, TRAM-34 did not evoke afferent discharge (main effect of drug, p = 0.55, Figure 5A). The peak change in afferent firing following TRAM-34 application (−1.8±0.52 spikes s^−1^, N = 6) was no different to that in vehicle control experiments (p = 0.45, Figure 5B).

**Figure 5:**
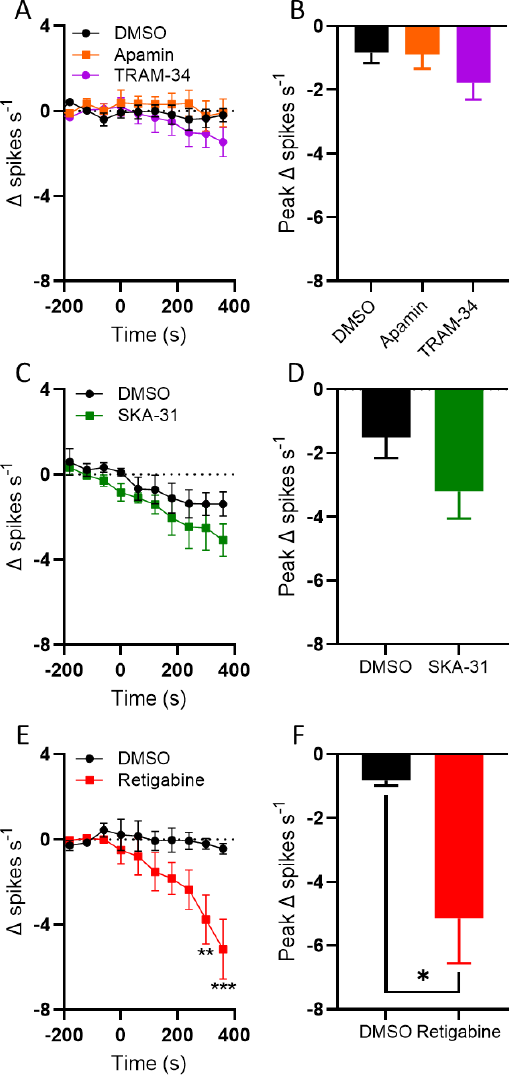
SK/IK channels do not regulate spontaneous colonic afferent activity. (A) Grouped data showing the change in spontaneous afferent firing rate following the application of DMSO, apamin (orange) or TRAM-34 (purple) (applied at 0 s). Two-way ANOVA. (B) Grouped data showing the peak change in afferent firing rate following DMSO, apamin or TRAM-34 application. One-way ANOVA with Holm-Sidak’s post-tests. (C) Grouped data showing the change in spontaneous afferent firing rate following the application of DMSO or SKA-31 (green). Two-way ANOVA. (D) Grouped data showing the peak change in afferent firing rate following DMSO or SKA-31 application. Two-tailed unpaired t-test. (E) Grouped data showing the change in spontaneous afferent firing rate following the application of DMSO or retigabine (applied at 0 s). Two-way ANOVA with Holm-Sidak post-tests. (F) Grouped data showing the peak change in afferent firing rate following the application of DMSO or retigabine. Two-tailed unpaired t-test.

Consistent with these findings, and those from earlier experiments, SKA-31 application had no effect on spontaneous nerve activity compared to DMSO-treated tissue (main effect of drug, p = 0.22, Figure 5C). The peak decrease in afferent firing rate after SKA-31 application (3.2±0.7 spikes s^−1^, N = 6) was no different to that after DMSO application (1.5±0.6 spikes s^−1^, N = 5, p = 0.16, Figure 5D).

Conversely, the application of retigabine suppressed spontaneous afferent firing ≥5 minutes after application (main effect of drug, p = 0.069; 5 min, p = 0.0052; 6 min, p < 0.0001, Figure 5E). The peak decrease in spontaneous afferent discharge in control experiments was 0.83±0.16 spikes s^−1^ (N = 5) compared to 5.2±1.4 spikes s^−1^ (N = 6) after retigabine application (p = 0.027, Figure 5F).

### SK/IK channels regulate colonic motility

Finally, to confirm the activity of SKA-31, we demonstrated its ability to inhibit ongoing peristaltic gut motility consistent with the known role of IK channels in mediating the afterhyperpolarisation of Dogiel type II enteric neurons. Pressure changes evoked by colonic migrating motor complexes (CMMCs) were recorded before and after the application of SKA-31 (100 µM, Figure 6A). CMMCs exhibited an average amplitude 16.6±6.9 mmHg before SKA-31 application and 5.1±2.2 mmHg after SKA-31 application (p = 0.0092, N = 4, Figure 6B). The frequency of CMMCs was also reduced from 0.35±0.03 min^−1^ to 0.15±0.03 min^−1^ by SKA-31 application (p = 0.045, N = 4, Figure 6C).

**Figure 6:**
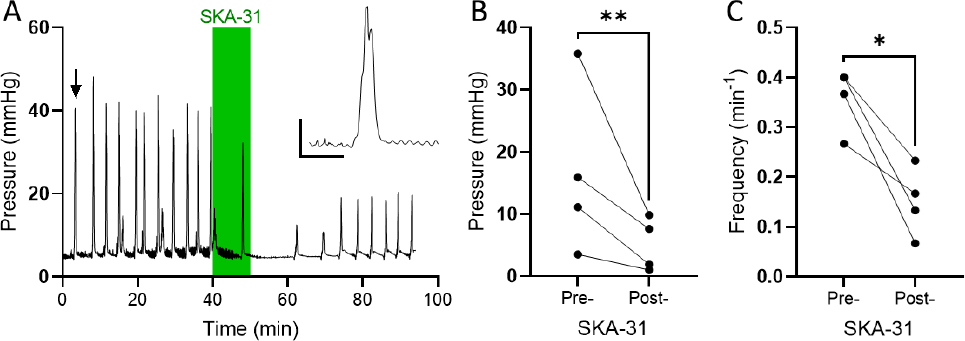
SK/IK channels regulate colonic motility. (A) Exemplar recording of colonic luminal pressure, showing the effect of SKA-31 application (green shaded area). *Inset*: expanded trace showing an individual CMMC (highlighted by the arrow). Scale: 1 min; 10 mmHg. (B) Grouped data showing the amplitude of CMMCs before and after the application of SKA-31. Two-tailed paired t-test. (C) Grouped data showing the frequency of CMMCs before and after the application of SKA-31. Two-tailed unpaired t-test.

## Discussion

We have investigated the role of SK and IK channels in regulating the colonic afferent response to ATP, bradykinin and distention of the colon. ATP and bradykinin are important algogenic stimuli known to contribute to visceral nociception and pain during inflammation. Distention of the colon is an important visceral stimulus in both physiological and pathological states. The colonic afferent response to all of these stimuli, as well as basal afferent activity, was unaffected by the opening of SK/IK channels by SKA-31.

This observation was unexpected given the expression of transcripts encoding SK channel subunits in back-labelled colonic afferent neuron cell bodies (Hockley *et al*., 2019). However, the expression data used here does not give any indication of protein expression at the afferent terminals in the colon. It may be that the cell bodies and terminals of colonic afferents express different subsets of ion channels, and that SK channels are only expressed at the cell body. If SK channels are expressed at the terminals of colonic afferent nerves, why did SKA-31 have no effect on afferent activity? It is possible that SK channels are tonically active under basal conditions (Liu and Herbison, 2008; Zhang and Huang, 2017), rendering SKA-31 unable to potentiate their activity any further. However, inhibition of SK channels with apamin did not enhance afferent activity, suggesting that SK channels are unlikely to be tonically active. The inhibition of IK channels with TRAM-34 also yielded no change in colonic afferent activity, indicating that IK channels are not likely tonically active either.

It is also possible for SK channel subunits form heteromeric channels with other members of the SK family, which can affect trafficking of channels to the membrane (Monaghan *et al*., 2004) and channel pharmacology (Benton *et al*., 2003; Monaghan *et al*., 2004; Church *et al*., 2015). For example, channels formed of SK1 and SK2 subunits exhibited an apamin sensitivity intermediate between SK1 and SK2 (Benton *et al*., 2003; Church *et al*., 2015). The high concentration of apamin used in the present study would still be expected to robustly inhibit heteromeric channels. Therefore, it is doubtful that the lack of effect of apamin observed here is result of the formation of heteromeric SK channels. SK1 subunits preferentially co-assemble with IK (rather than forming homomeric channels), forming SK1-IK channels insensitive to apamin and with a markedly reduced sensitivity to TRAM-34 compared to homomeric IK channels (Higham *et al*., 2019). ∼80% of *Kcnn4*-expressing colonic afferents in mouse co-express *Kcnn1*, indicating the potential for the presence of a significant population of heteromeric SK1-IK channels. Despite this, if functional, tonically active SK/IK channels were expressed at the afferent terminals in the manner suggested by expression at the cell bodies (Hockley *et al*., 2019), it would still be expected that colonic afferent activity could be elevated by apamin or TRAM-34. This is because *(i)* a large population of *Kcnn1*-expressing afferents do not express *Kcnn4*, and *(ii)* 10 µM TRAM-34 (used in this study) blocks ∼50% of heteromeric SK1-IK channel current (Higham *et al*., 2019), which would be expected to induce membrane depolarisation sufficient to trigger action potential discharge. Consequently, it seems unlikely that heteromeric channel formation underpins the lack of effect of apamin and TRAM-34 on colonic afferent activity. The effect of SKA-31 on heteromeric SK/IK channels is not known.

In all, it seems probable that the lack of effect of SKA-31, apamin and TRAM-34 is due to a lack of SK and IK channel expression on colonic afferent terminals. SK channels expressed in sensory neuron cell bodies could still control visceral sensory input to the central nervous system (Hao *et al*., 2023), but they do not appear to contribute to the control of colonic afferent activity at the terminals.

At odds with results obtained with SKA-31, we found that retigabine inhibited colonic afferent activity in response to stimulation with ATP, bradykinin and distention of the colon. Transcripts encoding receptors for ATP and bradykinin are frequently co-expressed with transcripts encoding K_V_7 channels subunits (*KCNQ2*, *KCNQ3* and *KCNQ5*) (Peiris *et al*., 2017; Hockley *et al*., 2019). Both ATP and bradykinin evoke action potential firing through multiple mechanisms, including the inhibition of K_V_7 channels in a phospholipase C-(PLC) dependent manner (Stemkowski *et al*., 2002; Liu *et al*., 2010). It is likely that many PLC-coupled receptors, such as protease activated receptor 2 (PAR2) (Linley *et al*., 2008) and Mas-related G-protein-coupled receptor member D (MrgD) (Crozier *et al*., 2007), stimulate action potential discharge through the inhibition of K_V_7 channels. Suppression of K_V_7 channel activity triggers membrane depolarisation necessary for the activation of voltage-gated Na^+^ (Na_V_) channels required to further amplify depolarisations (e.g., Na_V_1.9) and trigger action potential generation (e.g., Na_V_1.8). Indeed, nocifensive behaviours evoked by bradykinin injection and colonic afferent discharge evoked by ATP can both be markedly attenuated by genetic loss of Na_V_1.9 (Liu *et al*., 2010; Hockley *et al*., 2016). It seems clear that K_V_7 channels are an important regulator of the colonic afferent response to chemical stimuli.

Retigabine also robustly inhibited the afferent response to distention of the colon across a wide range of distention pressures (>5 mmHg). This indicates that K_V_7 channels likely regulate the function of low-threshold, wide dynamic range and high-threshold mechanosensitive afferents, in agreement with the broad expression of transcripts encoding the subunits comprising K_V_7 channels (Peiris *et al*., 2017; Hockley *et al*., 2019). It has previously been shown that retigabine suppressed mechanically-evoked activity in the saphenous nerve (Roza and Lopez-Garcia, 2008) and that linopiridine, a K_V_7 channel blocker, potentiated the activity of rapidly-adapting mechanoreceptors in the skin (Heidenreich *et al*., 2012).

Finally, we observed a robust suppression of the amplitude and frequency of CMMCs following the application of SKA-31, consistent with the well-established role for IK channels in the regulation of the excitability of Dogiel type II neurons (Mao, Wang and Kunze, 2006) which initiate peristaltic reflexes. Opening of SK or IK channels causes membrane hyperpolarisation, suppressing voltage-gated Ca^2+^ channel activity and, hence, reduces the Ca^2+^ influx required for smooth muscle contraction. SK channel expression has already been identified in colonic smooth muscle (Klemm and Lang, 2002). The tension of colonic muscle was increased by SK channel blockade, and oestrogen-induced colonic smooth muscle relaxation may be due, in part, to the opening of SK channels (Tang *et al*., 2015).

In summary, data presented here do not support the hypothesis that SK/IK channels regulate colonic afferent activity, though these channels do appear to regulate colonic smooth muscle activity. Our data do, however, provide further support for K_V_7 channels playing a key role in regulating the colonic afferent response to chemical and mechanical stimuli.

## Additional information

### Author contributions

Conception and design: CNB, TR and DCB. Performed experiments: CNB and RG. Analysed data: CNB and JPH. Wrote the manuscript: JPH and DCB. Acquired funding: TR and DCB.

## Acknowledgements

This work was supported by a Crohn’s and Colitis U.K. grant to D.C. Bulmer (PC2019/1-Bulmer). C.N. Bhebhe was supported by Gates Cambridge Trust scholarships.

